# An inducible and reversible system to regulate unsaturated fatty acid biosynthesis in *C. elegans*

**DOI:** 10.1101/2024.10.01.616130

**Authors:** Bernabe Battista, Bruno Hernadez Cravero, Monica P Colaiácovo, Luisa Cochella, Andres Binolfi, Diego de Mendoza

## Abstract

Unsaturated fatty acids (UFAs) play crucial roles in various physiological and pathological processes. In animals, these lipids are synthesized from saturated fatty acids through the action of delta 9 (Δ9) desaturases. In *C. elegans*, three Δ9 desaturases are encoded by the genes *fat-5*, *fat-6*, and *fat-7*. The presence of multiple Δ9 desaturases has posed a significant challenge in developing a rapid and efficient approach to control UFA production in *C. elegans* and other model organisms. Utilizing the auxin-inducible degradation system, we specifically targeted the *C. elegans fat-7* gene, responsible for the major stearoyl-CoA desaturase (SCD), while deleting *fat-5* and *fat-6*. This design resulted in a strain that can be reversibly depleted of UFAs in the cells of interest. Conditional depletion in all somatic cells exhibited a pronounced auxin-dependent defect in UFA production. Using this system, we uncovered an essential requirement for de novo UFA production during L1 and L2 stage. Moreover, our results support a direct connection between UFA levels, fat storage and increased lipid turnover. This system will enable further studies exploring the cellular and physiological consequences of impairing UFA biosynthesis at different developmental stages or in specific tissues.

**Summary:** Unsaturated fatty acids (UFAs) are essential for life. In animals, UFAs are synthesized by Δ9 desaturase enzymes. *Caenorhabditis elegans* possesses three Δ9 desaturase genes: fat-5, fat-6, and fat-7. We engineered a strain where fat-7 can be reversibly switched off while fat-5 and fat-6 are deleted, allowing precise control of UFA levels throughout the life cycle. Our findings demonstrate the critical role of UFA biosynthesis in early development and its direct link to fat storage and lipid turnover. This strain enables the study of UFA-related physiological and pathological processes in animals.

## Introduction

Lipids are key biologic molecules that contribute to a variety of cellular and organismal functions in three main manners (Watts and Ristow 2017). First, they are fundamental structural elements of cellular membranes, which build a selective barrier separating the cell from the environment and ensuring subcellular compartmentalization (Harayama and Riezman 2018). Second, they are key molecules in energy metabolism that fuel the cell (Watts and Ristow 2017; Yao et al. 2020). Third, they play active roles in signal transduction by either directly acting as signal molecules, or indirectly by affecting membrane fluidity (Kutchai et al. 1976; Kniazeva et al. 2004; Jeong et al. 2005; Zhu et al. 2013; Svensk et al. 2016; Busayavalasa et al. 2020). Thus, knowledge of the consequences of changes in lipid levels, composition and location has implications for understanding organismal biology in health and disease. A powerful animal model for trying to answer this question is the nematode *Caenorhabditis elegans*.

In practically all eukaryotic organisms, including *C. elegans*, unsaturated fatty acids (UFAs) are essential components of membrane and storage lipids. UFA synthesis depends on the conversion of saturated fatty acids by Δ9 desaturases (Brock et al. 2006; Brock et al. 2007). In *C. elegans* the *fat-6* and *fat-7* genes encodes a Δ9 stearoyl-CoA desaturase and a similar gene, *fat5* encodes a Δ9 palmitoyl-CoA desaturase. The pathway for unsaturated fatty acid (UFA) synthesis in *C. elegans* begins with palmitic acid (16:0), obtained from the *E. coli* diet or synthesized *de novo*, which is converted to palmitoleic acid (16:1 Δ9) by FAT-5. This fatty acid is then elongated to cis-vaccenic (18:1Δ11), which is an abundant fatty acid in phospholipids and triglycerides. Palmitic acid (16:0) can also be elongated to stearic acid (SA, 18:0), the substrate for FAT-6 and FAT-7 desaturation to oleic acid (OA, 18:1Δ9) (Figure 1 and Figure S1). Unlike most animals, *C. elegans* possesses a Δ12 fatty acid desaturase, which allows it to synthesize all polyunsaturated fatty acids (PUFAs) from OA (Figure S1)(Brock et al. 2007; Watts and Ristow 2017). These PUFAs play crucial roles in membrane function and signaling (Watts 2016). Previous genetic and phenotypic studies showed that the three Δ9 desaturases single mutants, *fat5*, *fat6* and *fat7* display few differences from wild type because they compensate for loss of one isoform by regulated induction of the remaining Δ9 desaturase genes (Brock et al. 2006; Brock et al. 2007). However, the Δ9 desaturation activity is essential, and *fat6; fat7 fat5* triple mutants that lack this activity are unable to survive unless they are supplemented with an exogenous fatty acid mix composed of monounsaturated fatty acids (MUFAs) and polyunsaturated fatty acids (PUFAs)(Brock et al. 2006).

**Figure 1.**
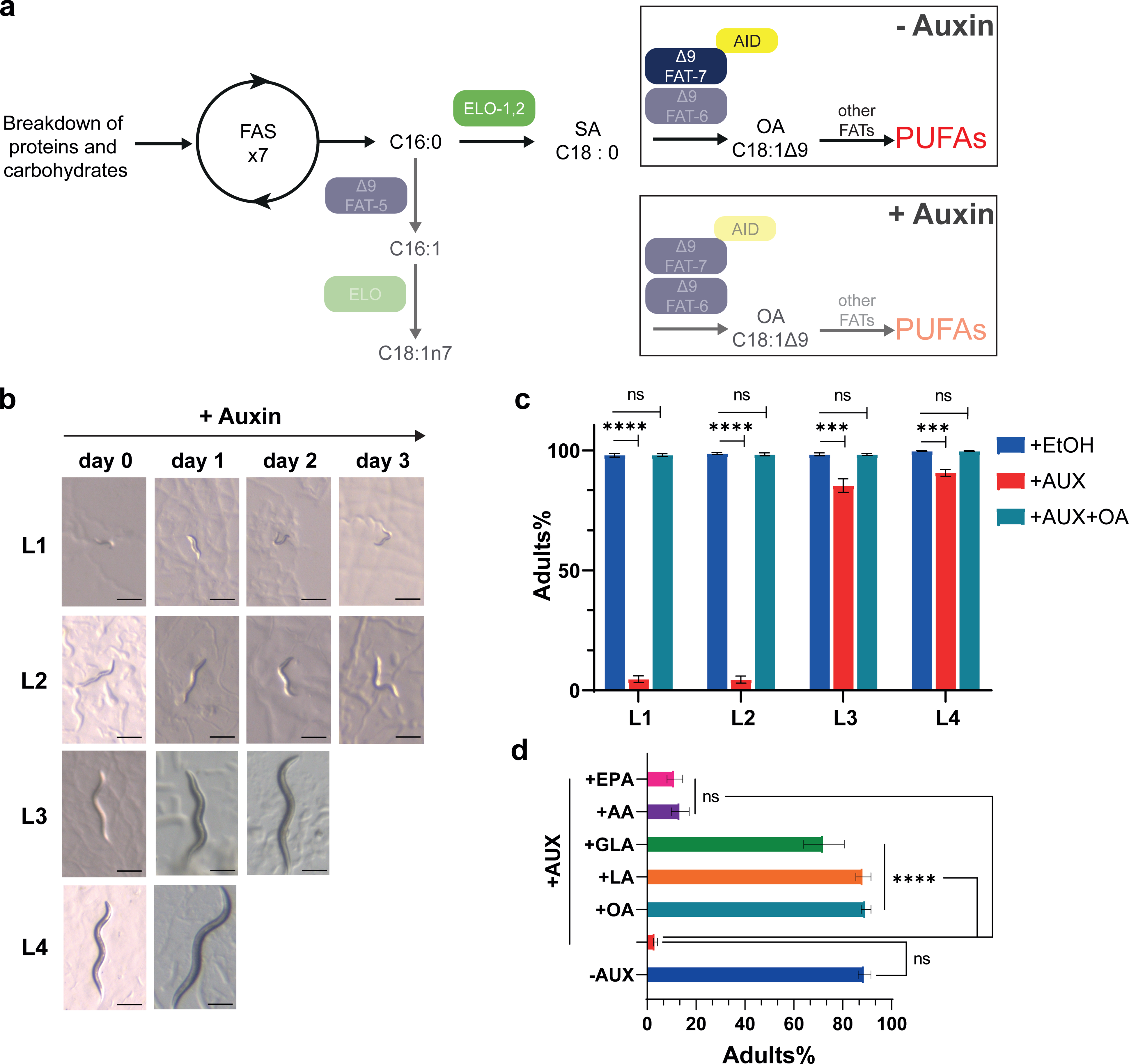
Auxin-induced interruption of UFA synthesis leads to developmental arrest in DDM6 worms. **(a)** Simplified *de novo* fatty acid synthesis pathway of *C. elegans* DDM6 mutant strain in presence and absence of auxin. Enzyme names and activities are enclosed in ovals. FAS, fatty acid synthase; 16:0, palmitic acid, 16:1, palmitoleic acid, 18:1n7, cis-vaccenic acid; 18:1Δ9, oleic acid; PUFAs, polyunsaturated fatty acids. **(b)** Representative images of L1, L2, L3 and L4 larvae growing on auxin supplemented NGM plates for several days. Scale bar = 200 µm. **(c)** Average percentage of adults reached by DDM6 L1, L2, L3 or L4 larvae incubated on auxin, ethanol, or auxin + fatty acid-supplemented plates at 20°C. Values significantly different from DDM6 + ethanol worms using unpaired *t*-test are (*) p < 0.05; (**) p < 0.01; (***) p < 0.001 and (****) p < 0.0001. Data shown are average of 3 independent experiments, each of three biological replicates; error bars indicate SEM. **(d)** Average percentage of adults reached by DDM6 L1 larvae incubated on auxin with and without fatty acid supplementation. Values significantly different from DDM6 + auxin worms using non-parametric Kruskal-Wallis test followed by Dunn’s multiple comparisons correction test, are (*) p < 0.05; (**) p < 0.01; (***) p < 0.001 and (****) p < 0.0001. Data shown are average of 4 to 9 independent experiments, each of three biological replicates; error bars indicate SEM.

Given that UFAs play pivotal roles in membrane biology, efforts have been undertaken in *C. elegans* to disrupt the synthesis of these fatty acids, aiming to investigate the cellular and physiological consequences of compromised membrane homeostasis. A recently employed strategy for diminishing UFA synthesis involves utilizing *paqr-2*, *paqr-1* double mutants (Svensson et al. 2011; Busayavalasa et al. 2020). PAQR-2 and PAQR-1 exhibit partially redundant functions in promoting fatty acid desaturation and the integration of UFAs into phospholipids (Svensk et al. 2013; Svensk et al. 2016; Busayavalasa et al. 2020; Devkota, Henricsson, et al. 2021). However, recent studies have demonstrated that in addition, PAQR-2 amplifies sphingosine 1-P (S1P) production in response to membrane rigidification. S1P, a vital bioactive lipid, actively participates in diverse cellular and physiological processes, including binding to transcription factors that boost the transcription of *fat-5*, *fat-6*, and *fat-7* in *C. elegans* (Svensk et al. 2013; Vasiliauskaité-Brooks et al. 2017; Pilon 2021; Ruiz et al. 2022). Consequently, *paqr* mutants display high pleiotropy, potentially complicating the interpretation of the physiological consequences of UFA deprivation (Devkota, Kaper, et al. 2021). As a result, the field of membrane lipids has thus far lacked a genetically defined animal model to selectively obstruct UFA biosynthesis. Such a model would prove invaluable in advancing our comprehension of the molecular underpinnings of numerous processes affected by UFAs.

**(a)** *C. elegans* lacks fat storage cells and stores neutral lipids, mainly triglycerides (TAG) and low amounts of cholesterol and cholesterol esters, in intestinal and hypodermal lipid droplets (LDs) (Mak 2012). LDs supply fatty acids to β−oxidation processes for energy production and serve as shuttles for lipids and proteins among different tissue and sub-cellular compartments (Pol et al. 2014). *C. elegans* strains with impaired Δ9 stearoyl-CoA desaturase have reduced adiposity and LD size (Shi et al. 2013). Moreover, recent work showed that *C. elegans* PUFAs stimulate LD fusion and growth (Wang et al. 2022), and that FAT-7 is directly involved in regulating organismal lipid storage and LD dynamics in an S-adenosyl-methionine and phosphatidylcholine dependent manner (Han et al. 2024). Thus, a conditional mutant where the total UFA synthesis is blocked at selected developmental stages represents a powerful tool to understand the role of UFAs in lipid storage biology and metabolic syndromes, such as obesity (Xie et al. 2022).

Here, we used the auxin-inducible degradation (AID) system in *C. elegans* (Zhang et al., 2015) to create a strain in which *fat-7* can be conditionally depleted. When combined with a double mutant of *fat-5* and *fat-6*, this setup allows for the targeted, acute depletion of OA in cells that express the TIR1 auxin-dependent, ubiquitin ligase. We show that expression of TIR1 in somatic cells enables auxin-dependent conditional depletion of MUFA synthesis at different larval stages. Our characterization of the conditional mutant reveals an essential role for *de novo* synthesis of OA in the transition from L1 and L2 larvae to the adult stage. We also demonstrate that auxin-mediated L1 larval arrest is caused by the inability to synthesize OA (18:1Δ9) or C18 diunsaturated fatty acids, rather than by a lack of C20 PUFAs. Moreover, depleting FAT-7 at the L3 stage interferes with lipid storage by decreasing the size and TAG levels in LDs, a phenotype that is efficiently rescued by exogenous supplementation of OA. Finally, we show that exposure to auxin at the L3 stage induces sterility as shown by a marked decrease in hermaphrodite self-progeny, which can be reversed upon OA media supplementation. Together, our results demonstrate that conditional depletion of tagged FAT-7 in the absence of redundant enzymatic activities provides a powerful new tool for the spatiotemporal regulation and analysis of UFA function in a metazoan model organism.

## Materials and Methods

### Nematode maintenance and strains

Unless otherwise noted, worms were grown on nematode growth media plates (NGM) at 20°C seeded with E. coli OP50 lawns as a source of food (Kenyon 1988). Bacterial cultures were grown overnight on LB-broth (BD-Difco). Auxin solution was prepared in ethanol as carrier, then it was added to seeded NGM plates to a final concentration of 4mM along with hexane. Fatty acid methyl-ester were prepared in hexane from commercial pure fatty acid (see below) and were used to supplement the auxin plates to a final concentration of 200 µM. Control plates were prepared by adding the same amounts of ethanol and hexane as in the auxin plates to seeded NGM plates. All plates were freshly prepared and vented for about 1.5-2 hours to allow full solvent evaporation, then stored at 4°C protected from light and used within a week. N2 Bristol was employed as wild-type strain. The other strain used in this research was DDM6 [ieSi57 II; unc-119(ed3) III; fat-6(tm331) IV; fat-7(dme1[fat-7::3xflag::aid]) fat-5(tm420) V].

### Construction of fat-7::3Xflag::AID and generation of DDM6 strain by CRISPR/Cas9 genome editing

To generate a C-terminal AID-tagged fat-7, CA1200 [ieSi57(Peft-3::TIR1::mRuby)] animals were subjected to Cas9 mediated genome engineering via RNP-microinjection following the Co-CRISPR protocol with modifications (Kim et al. 2014; Dickinson and Goldstein 2016). See supplemental material for a more detailed description.

Once fat-7(dme1[fat-7::3xflag::aid]) was generated, it was crossed into strain BX110 carrying fat-6(tm331) IV; fat-5(tm420) V deletions to generate the UFA deficient conditional mutant strain DDM6: ieSi57 II; unc-119(ed3) III; fat-6(tm331) IV; fat-7(dme1[fat-7::3xflag::aid]) fat-5(tm420) V.

### Fatty acid extraction for GC-MS

For fatty acid analysis, c.a. 40000 N2 or DDM6 L3 larvae were transferred to auxin/auxin+OA/ethanol plates and allowed to grow for 48h (1-day old adults) or 72h (2-day old adults). Then, the adult worms were harvested and washed 3 times with M9 to remove any excess of bacteria. Worms were centrifuged, excess of buffer was removed, and the pellet was transferred to a glass tube. Lipids were extracted with 3 mL of chloroform:methanol 1:2 mixture (Blight-Dyer method). Dibutylhydroxytolueno (BHT) was added as an antioxidant at a final concentration of 0.005% v/v. The samples were incubated overnight at -20°C, centrifuged and the supernatants were transferred to a new tube. 1 mL of 1M KCl 0.2M H3PO4 and 1mL chloroform were added to the samples before vortex and centrifugation for 2 min at 2000g. Finally, the organic phase was transferred to a new tube and dried under N2(g) flow.

### Fatty acid methyl-ester preparations for GC-MS

For the derivatization of fatty acids previous to GC-MS analysis, 2mL of 2.5% H2SO4 in methanol were added to the lipid samples, the tubes were capped and incubated at 80°C for 2h. The fatty acid methyl-esters were extracted with 3 mL of a 1:1 mixture of hexane and 5% v/v NaCl, dried under N2(g) flow and, finally resuspended in 500 µL of hexane.

### Gas chromatography coupled to mass spectrometry (GC-MS)

The methyl ester analysis was performed in a Shimadzu GC-2010 Plus equipment. The column used was a Supelco WAX-10 (Sigma Aldrich) 100% polyethyleneglycol. The helium flux was 1 ml/min and the heating program was 180 °C (0 min to 32 min) and then an increasing gradient of 3 °C/min from 180 °C to 240 °C. The split was 1/30 and the ionization voltage was 70 eV with an ionic range from 50 to 600 Da. The ion specters were registered as relative abundance in function of mass/charge (m/z). Peak assignation was done using the mixture of standards of fatty acid methyl esters PUFA3 (Matreya).

### Sample preparation for in vivo NMR experiments

13C isotopically enriched worms for in vivo and lipid extracts experiments were prepared as previously reported (Hernández-Cravero et al. 2024). Briefly, 60000 synchronized DDM6 L1 larvae/sample were placed on NGM plates seeded with 20X pellet of an overnight culture of E. coli OP50 grown in a M9 minimal medium supplemented with 12C-D-glucose (Sigma) and 1 g/L of 14N-ammonium chloride (Merck) as sole carbon and nitrogen sources, respectively. When the worms reached the L3 stage, they were transferred to auxin/auxin+OA/ethanol NGM plates seeded with 20X pellet of an overnight culture of E. coli OP50 grown in a M9 minimal medium supplemented with 13C-D-glucose (Cortecnec, France) and 1 g/L of 14N-ammonium chloride (Sigma-Aldrich) as sole carbon and nitrogen sources, respectively. OA was not isotopically enriched. In all cases M9 media was also supplemented with 20mg/L uracil (Merck) to compensate for OP50 auxotrophy under this minimal media. Animals were grown until 1-day adulthood, collected in a 15 mL falcon tube, centrifuged for 2 min at 2000 g to remove media and washed twice on M9 buffer to remove remaining bacteria. Supernatant was discarded, and M9 buffer was added to complete a volume of 700 μL supplemented with 10% D2O (99.9%, Sigma). Afterwards, worms were loaded in a 5 mm NMR Shigemi tube (without applying the plunge) and decanted to the bottom by gentle spinning.

### Fatty acid extraction for in vitro NMR experiments

For fatty acid analysis of C. elegans extracts, nematodes were grown until 1-day adult stage, as for GC-MS. Then, nematodes were harvested and washed 3 times with M9 to remove any excess of bacteria, centrifugated, and pellets were frozen at -80°C. The pellets were thawed, resuspended in 1.3 mL of pure methanol, separated in 3 tubes and sonicated on ice (Bioruptor sonifier, Diagenode). To prevent samples from heating, periods of 30 s at maximum power sonication were alternated with sonication-free lapses of 1 min during 10 min. After sonication, 2.6 ml of chloroform and 1.3 mL 0.5 M KCl/0.08 M H3PO4 were added to a final ratio of 1:2:1. The solutions were first subjected to sonication in an ultrasonic water bath for 15 minutes, followed by 2 minutes of vortex mixing. They were then centrifuged at 2,000 g for 10 minutes to facilitate phase separation. The lower, hydrophobic phase was carefully transferred into a clean glass tube, dried under nitrogen stream, and re-dissolved in 500 μL of deuterated chloroform containing 0.005% BHT (Butylated hydroxytoluene) (Sigma) to prevent lipid oxidation. Finally, the solution was transferred into 5 mm NMR tubes (Norell, USA) using a glass pipette.

### NMR spectroscopy

NMR spectra were registered at 25 oC on a 700 MHz Bruker Avance III spectrometer and a 5 mm triple resonance TXI probe (1H/D 13C/15N). 1D 1H spectra of lipid extracts were registered with a 30-degree flip angle hard pulse sequence (zg30), 8K points, recycling delay of 3 s, 512 scans and spectral width of 16 ppm. Processing was done by zerofilling to 32k points followed by a qsine window function multiplication, Fourier Transform, phase and baseline correction. 1D 1H spectra of live worms and supernatant samples were acquired using a pulse sequence with excitation sculpting for water suppression (zgesgp) (Hwang and Shaka 1995), 8K points, a recycling delay of 1 s, 128 scans and spectral width of 16 ppm. Processing was done by zerofilling to 32k points following by a qsine window function multiplication, Fourier Transform, phase and baseline correction. 2D 1H-13C HSQC spectra of 13C-isotopically enriched lipid extracts were acquired with a phase-sensitive pulse sequence and gradient pulses (hsqcgpph). We used 1K and 256 points in 1H and 13C, respectively, 4 scans and 16 ppm and 160 ppm spectral width for 1H and 13C, respectively. 13C decoupling was done with the GARP sequence. The spectra were processed with qsine window function multiplications, Fourier Transform and phase and baseline correction in both dimensions. 2D 1H-13C HSQC spectra of live worms were acquired on 13C-isotopically enriched animals, with a sensitivity-enhanced pulse sequence (hsqcetgpsisp2.2). We used 2K and 256 points in 1H and 13C, respectively, 4 scans and 16 ppm and 150 ppm spectral width for 1H and 13C, respectively. Experimental time was 36 min. 13C decoupling was done with the GARP sequence. After NMR acquisitions, the worms were transferred to an eppendorf tube, and the supernatant was separated by gentle centrifugation (10 min at 1000 g).

### Growth analysis

Worms were synchronized by hypochlorite treatment (Porta-de-la-Riva et al. 2012). Eggs were allowed to hatch overnight in M9 buffer and L1 larvae were placed onto seeded NGM regular plates, so they continue their development. Synchronous populations of L1, L2, L3 and L4 were collected from those plates and transferred to auxin/ethanol/auxin+FAME and incubated at 20°C. The number of adult animals was scored within the following 3 days.

### Fertility analysis

To analyze the progeny of individual worms, L4 (grown on auxin/ethanol/auxin+OA from L3 stage as described before) were isolated and moved to auxin/ethanol/auxin+OA fresh plates. After reaching reproductive age, adults were moved daily to a fresh plate for 48h. The adult worms were removed before the total live progeny was counted. For the reversibility assays, L4 hermaphrodites (grown on auxin from the L3 stage as previously described) were isolated and transferred to auxin plates for 24 hours, then moved to fresh control plates (+EtOH) daily for 48 hours (condition +/-). In the +/+ condition, after the initial 24-hour exposure to auxin, worms were transferred to freshly prepared auxin plates daily for 48 hours. Adult worms were removed before counting the total number of live progeny.

### Survival at low temperatures

Synchronous populations of L3 were moved to auxin/ethanol/auxin+OA and incubated at 20°C, 15°C and 10°C for several days. The number of live, non-arrested worms was counted on each plate when the control population (+EtOH) reached adulthood. These values are expressed relative to the number of live non-arrested worms counted at 20°C (Brock et al. 2007).

### Nile red staining and quantification

Adult worms were fixed and stained with Nile red as described in (Nhan et al. 2019). For both total lipid droplet intensity and size quantification, worms were imaged using a Nikon Eclipse 800 equipped with a Andor Clara digital camara at 10X and 40X magnification, respectively. For total fluoresce intensity determination, intensity was relativized to whole worm area for each animal (Nhan et al. 2019). Total lipid droplet size determinations were made similarly as described in (Shi et al. 2013). Briefly, for each worm photograph, a 18x18 µm square was placed arbitrarily over the mid-intestinal region, and within that square each visible Nile Red-stained droplet was determine automatically using the ImageJ software. For each worm, the average lipid droplet size was determined. Statistical comparisons (Kruskal-Wallis and Dunn’s multiple comparisons test) were performed using GraphPad Prism 8.

## Results

### Depletion of UFAs induces phenotypic defects in *C. elegans*

Since a *fat-6; fat-7 fat-5* triple mutant is unable to survive (Brock et al. 2006), we exploited the *Arabidopsis* auxin-inducible degron (AID) tool kit system to engineer the *fat-7* locus using CRISPR/Cas9 to generate a strain for conditional depletion of FAT-7 (Figure S2). AID relies on the exogenous expression of the plant F-Box protein TIR1, which mediates the depletion of the degron tagged targets upon exposure to auxin (Zhang et al. 2015; Ashley et al. 2021)(see reagent table). We tagged endogenous *fat-7* with the TIR1 ubiquitin ligase-recognition peptide and expressed TIR1 ubiquitously in all somatic cells. Then, we combined these components with available *fat-6* and *fat-5* null alleles to generate a UFA deficient conditional mutant, hereafter referred to as DDM6 (Figure 1a). To test whether this strain produced the auxin-dependent conditional lethal phenotype expected for an animal lacking all three Δ9 desaturases, worms were transferred to NGM plates seeded with *E. coli* OP50 supplemented with auxin or ethanol (control plates) at various developmental stages, from L1 to L4 (Figure 1b and 1c). Auxin treatment starting at the L1 or L2 larval stage caused a completely penetrant arrest, in contrast to control animals that developed to adulthood. However, animals treated with auxin starting at the L3 or L4 stages were able to mature into adulthood. These observations uncover an essential requirement for de novo synthesis of UFAs during L1 and L2 and suggest that development of later stages relies on fatty acids synthesized prior to FAT-7 auxin-induced depletion (Figure 1b).

### Dietary supplementation of fatty acids suppresses the arrest phenotype depending on the number of atoms and degree of unsaturation

To investigate the potential reversal of the arrest phenotype and its specificity, we performed a dietary supplementation of different UFA methyl esters in L1 larvae treated with auxin (Figure 1d). Consistent with the expected biosynthesis block, the developmental arrest observed in L1 or L2 larvae exposed to auxin was completely reversed upon OA supplementation. Furthermore, the L1 arrest of DDM6 exposed to auxin was restored with exogenous addition of linoleic acid (LA, 18:2-n3) (Figure 1d), the product of OA desaturation by the FAT-2 (Δ12) desaturase (Figure S1). Partial rescue was also achieved with gamma-linoleic acid (GLA, 18:3n6) (Fig. 1D), the desaturation product of LA by the FAT-3 (Δ5) desaturase (Figure S1). In contrast, supplementation with C20 fatty acids from either the n-6 or n-3 series, such as arachidonic acid (AA, 20:4n6) and eicosapentaenoic acid (EPA, 20:5n3) respectively, did not reverse the arrest of L1 larvae induced by auxin exposure (Figure 1d). This indicates that these PUFAs are incapable of compensating for the deficiency of other UFAs synthesized by upstream desaturases (Figure S1). These findings align with previous observations suggesting that AA acid and EPA are non-essential for life in this organism (Vrablik and Watts 2013).

### *De novo* UFAs synthesis is required for growth at low temperatures and fertility

Previous studies have demonstrated that worms with impaired UFA synthesis exhibit sensitivity to cold stress (de Mendoza and Pilon 2019). To investigate whether shutting off UFA synthesis in DDM6 plays a role in survival at low temperatures, L3 worms grown at 20°C were transferred to plates containing auxin, ethanol (control), or auxin supplemented with OA and incubated at three different temperatures: 20°C, 15°C, and 10°C. We then scored the live, non-arrested worms after two days, which is the time it took control L3 worms to reach adulthood. Setting the survival rate at 100% at 20°C for control worms, we observed 100% and 79% survival at 15°C and 10°C, respectively (Figure 2a). These survival rates are consistent with those of the N2 strain under similar environmental conditions (Brock et al. 2006). However, worms placed on auxin plates showed a significant reduction in survival rates at low temperatures, with 81%, 27%, and 0% survival at 20°C, 15°C, and 10°C, respectively (Fig. 2A). In contrast, media supplementation with OA resulted in survival rates indistinguishable from those of the control group in all cases. Thus, interrupting *de novo* UFA biosynthesis at the L3 stage proves detrimental for nematodes at low temperatures.

**Figure 2.**
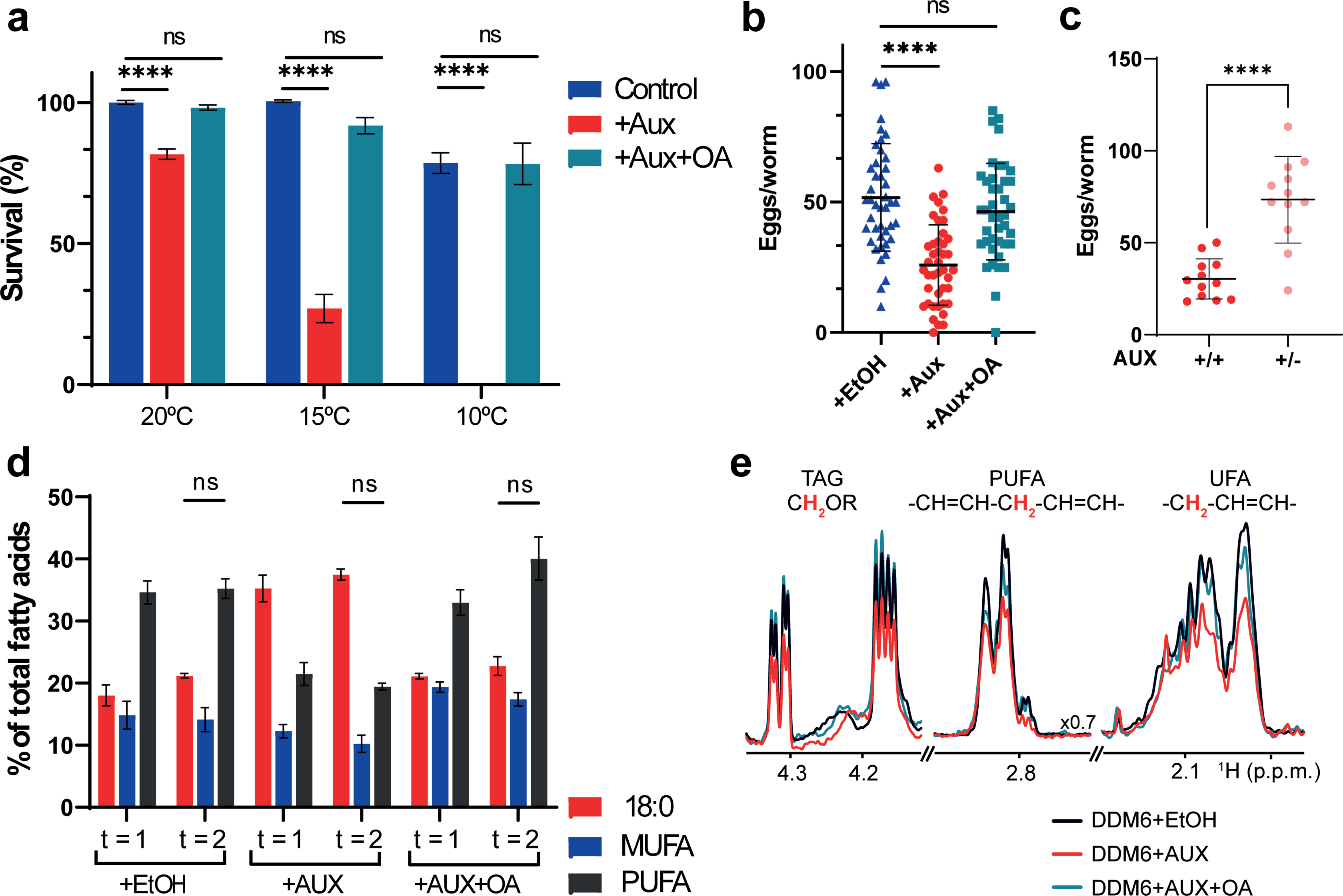
Reduced survival at low temperatures and fertility of DDM6 L3 larvae incubated with auxin. In all cases DDM6 L3 larvae were placed on auxin, ethanol or auxin + oleic acid supplemented NGM plates. **(a)** Survival at 15°C and 10°C relative to that of 20°C worms. Values significantly different from DDM6 + ethanol worms using unpaired *t*-test are (*) p < 0.05; (**) p < 0.01; (***) p < 0.001 and (****) p < 0.0001. Data shown are average of 2 independent experiments, each of six biological replicates; error bars indicate SEM. **(b)** Average number of progeny produced by individual adults during a 48h period. Values significantly different from DDM6 + ethanol worms using non-parametric Kruskal-Wallis test followed by Dunn’s multiple comparisons correction test, are (*) p < 0.05; (**) p < 0.01; (***) p < 0.001 and (****) p < 0.0001, n = 40 individuals / condition, error bars indicate SD. **(c)** Average number of progeny produced by individual adults during a 72h period. In +/+ condition, worms were exposed to auxin at all times, while in +/- condition worms were exposed to auxin for a 24h window and then changed to control plates for 48h hours. Values significantly different from +/+ worms using Welch’s t-test are (****) p < 0.0001, n = 12 individuals / condition, error bars indicate SD **(d)** Simplified fatty acid composition of one-day-old (*t* = 1) and two-day-old (*t* = 2) worms. 18:0, stearic acid; MUFA, total monounsaturated fatty acids, PUFA, total polyunsaturated fatty acids. Values significantly different from one-day-old worms of the same condition using unpaired *t*-test are (*) p < 0.05; (**) p < 0.01; (***) p < 0.001 and (****) p < 0.0001. Data shown are average of four to five independent determinations; error bars indicate SEM. **(e)** 1D ^1^H NMR of *C. elegans* lipid extracts in the TAG, PUFA and UFA spectral regions. The position of these regions in the spectra is indicated in Figure S3a (dotted squares). Extracts were prepared from DDM6 worms treated with ethanol (black), auxin (red), and auxin and oleic acid (cyan).

Even at the permissive temperature, (20^0^C), we observed a reduced brood size in DDM6 worms upon FAT-7 depletion compared to the control (Figure 2b). By counting the number of embryos produced by single worms during a 48 hour period, we found that DDM6 treated with auxin produced 50% of the progeny of the control condition (DDM6+EtOH). Supplementation with OA restored the progeny rate to control levels in auxin-treated DDM6 worms (Figure 2b), consistent with previous work showing that mutants impaired in PUFA synthesis have decreased lipid mobilization from intestine to oocytes, affecting early oogenesis and reducing the brood size (Lynn et al. 2015; Devkota, Kaper, et al. 2021). We also found that the reduction in brood size induced by auxin is reversible. When hermaphrodites exposed to auxin for a 24-hour window were transferred to control plates (+EtOH), the number of embryos deposited on the media increased (Figure 2c).

### Depletion of OA at the L3 stage alters fatty acid composition

To characterize the changes in lipid composition associated with the conditional depletion of OA, we transferred L3 larvae to auxin-containing plates until they reached the one-day adult stage and then profiled fatty acids using GC/MS (Table 1). The DDM6 mutant, when supplemented with auxin, showed significant changes in fatty acid composition compared to the control worms, with a notable accumulation of stearic acid (SA, 18:0). SA comprised 35.3% of the total fatty acids, a marked increase compared to 18% in the control worms. Consistent with this, there was a marked reduction in OA levels and its derived PUFAs, such as eicosatetraenoic acid (ETA, 20:4n3) and EPA, following auxin-induced FAT-7 degradation. Supplementing with OA largely restored the fatty acid composition of auxin-treated worms to levels comparable to those of the control group (Table 1), confirming that auxin-mediated depletion of this MUFA alters PUFA production, which is responsible for the associated physiological defects.

**Table 1.** Data are weight percentages (Mean+SEM) of four to six independent determinations of total worm fatty acids measured by gas chromatography. For each determination L3 larvae were placed on auxin, ethanol or auxin + oleic acid supplemented NGM plates until they reached the 1-day-old adult stage. 17Δ, 9,19-methylenehexadecanoic acid, 19Δ, 11,12-methyleneoctadecanoic acid. Values determined to be significantly different from DDM6+EtOH worms using an unpaired *t*-test are (*) p < 0.05; (**) p < 0.01; (***) p < 0.001 and (****) p < 0.0001.

| Fatty Acid | DDM6+EtOH | DDM6+Aux | DDM6+AUX+OA | N2+EtOH |
| --- | --- | --- | --- | --- |
| 14:0 n | 1.9 ± 0.7 | 1.8 ± 0.7 | 0.9 ± 0.09 | 2.0 ± 0.5 |
| 16:0 n | 6.1 ± 1.7 | 7.8 ± 1.1 | 5.3 ± 0.4 | 6.0 ± 1.4 |
| 18:0 n | 18.0 ± 2.9 | 35.3 ± 3.7 * | 21.1 ± 0.7 | 13.3 ± 2.5 |
| Total saturated | 26.1 ± 5.2 | 44.9 ± 5.2 | 27.4 ± 1.1 | 21.3 ± 4.1 |
| 16:1 cis | 0.7 ± 0.3 | 0.9 ± 0.3 | 0.6 ± 0.03 | 1.4 ± 0.4 |
| 18:1Δ9 | 4.0 ± 0.2 | 1.0 ± 0.1 *** | 9.2 ± 0.9 ** | 3.0 ± 0.6 |
| 18:1Δ11 | 10.0 ± 3.6 | 10.5 ± 1.7 | 9.5 ± 0.5 | 10.2 ± 2.3 |
| Total MUFA | 14.8 ± 3.9 | 12.3 ± 1.9 | 19.4 ± 1.4 | 14.7 ± 2.7 |
| 18:2 n6 | 11.6 ± 2.1 | 9.7 ± 2.5 | 12.7 ± 1.6 | 11.2 ± 1.1 |
| 20:3 n6 | 1.8 ± 0.3 | 1.4 ± 0.6 | 1.4 ± 0.02 | 2.8 ± 1.0 |
| 20:4 n6 | 1.8 ± 0.3 | 1.4 ± 0.6 | 1.4 ± 0.02 | 1.2 ± 0.2 |
| 20:4 n3 | 4.3 ± 0.2 | 2.4 ± 0.4 * | 4.2 ± 0.4 | 5.0 ± 1.1 |
| 20:5 n3 | 13.5 ± 1.5 | 5.8 ± 0.2 ** | 12.4 ± 0.8 | 11.7 ± 2.2 |
| Total PUFA | 34.6 ± 3.2 | 21.5 ± 3.1 * | 32.9 ± 3.5 | 32.0 ± 4.7 |
| Total UFA | 49 ± 6.7 | 33.7 ± 4.9 * | 52.3 ± 2.1 | 46.8 ± 7.2 |
| C15 iso | 6.3 ± 1.2 | 5.1 ± 0.9 | 3.6 ± 0.3 | 8.2 ± 1.3 |
| C17 iso | 8.5 ± 1.6 | 9.0 ± 1.5 | 4.6 ± 0.6 | 14.5 ± 1.5 * |
| Total branched | 14.8 ± 2.8 | 14.1 ± 1.8 | 8.2 ± 0.9 | 22.7 ± 2.8 |
| 17Δ | 6.0 ± 3.5 | 4.5 ± 1.1 | 10.2 ± 1.3 | 3.3 ± 1.1 |
| 19Δ | 3.5 ± 2.8 | 2.6 ± 0.3 | 1.7 ± 0.4 | 5.8 ± 3.0 |
| Total cyclopropane | 9.5 ± 2.7 | 7.1 ± 1.4 | 11.9 ± 1.3 | 9.1 ± 2.2 |

Independent ^1^H NMR experiments on extracts of unmodified lipids confirmed a decrease in the total contents of UFAs and PUFAs in DDM6 worms treated with auxin. We observed a reduction of ∼50% in the NMR signal amplitude of UFA and PUFA signals (Figure 2e and S3a) compared with control worms. Additionally, auxin treatment resulted in a similar reduction of triacylglycerol (TAG) molecules, as indicated by the lower intensity of the NMR signals corresponding to the - CH_2_OR-glycerol head group. The total levels of UFA, PUFA, and TAG were restored with the exogenous addition of oleic acid (OA), suggesting that FAT-7 activity is essential for TAG biosynthesis and/or increased storage to some extent.

We found that longer incubation with auxin does not induce further alterations in the fatty acid composition (Figure 2d, Table S1). Thus, the SA, total MUFA, and total PUFA levels of 1-day-old adults exhibited only subtle differences compared to 2-day-old adult worms, when both were incubated with auxin starting at the L3 larval stage.

### The interruption of UFAs biosynthesis reduces adiposity in DDM6 worms

We observed that DDM6 worms incubated with auxin often exhibit a clear intestine, a phenotype associated with reduced adiposity (Chen et al. 2019). To determine if this was the case, we used the lipophilic dye Nile Red to stain fat deposits and lipid droplets (LDs) in DDM6 animals whose UFA synthesis was interrupted at the L3 stage. Quantification of the total fluorescence confirmed that these worms had reduced fat storage compared to control worms, with about 60% of the total fluorescence intensity of the control group (Figure 3a, b).

**Figure 3.**
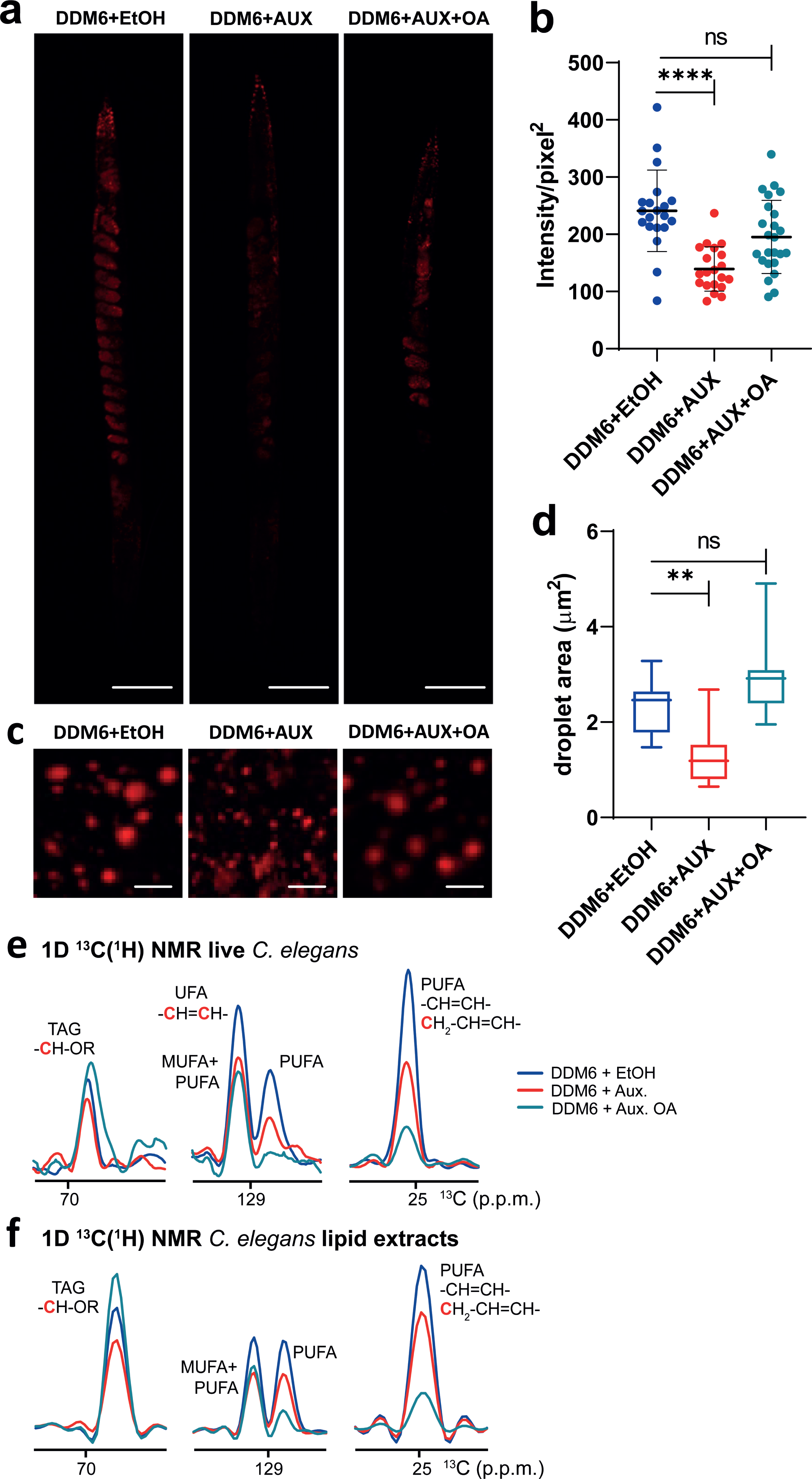
UFAs synthesis interruption alters lipid storage. **(a)** Fluorescent images of representative adult worms stained with Nile Red. Scale bars = 50 µm. **(b)** Nile red staining quantification, each dot represents the mean intensity value corrected by area for every worm scored, n = 20, error bars represent SD. **(c)** Fluorescent images of lipid droplets in the mid intestinal region showing a range of droplets size. Scale bars = 5 µm. **(d)** Quantification of lipid droplet size in the mid intestinal region. For **(b)** and **(d)** values significantly different from DDM6 + ethanol worms using non-parametric Kruskal-Wallis test followed by Dunn’s multiple comparisons correction test, are (*) p < 0.05; (**) p < 0.01; (***) p < 0.001 and (****) p < 0.0001. Data shown are average of 2 independent experiments. **(e, f)** ^13^C NMR spectra of live worms **(e)** and total lipid extracts **(f)** in the TAG, UFA and PUFA regions. DDM6 worms were isotopically enriched with ^13^C and treated with ethanol (blue), auxin (red) and auxin and non-isotopically enriched, ^12^C-oleic acid (cyan). ^13^C NMR traces were extracted from the 2D ^1^H-^13^C HSQC spectra shown in supplementary figure 4.

In addition, DDM6 worms exposed to auxin had smaller LDs compared to control worms (Figure 3c). The lipid droplet area in control worms ranged from 1.47 µm² to 3.28 µm², with an average size of 2.46 µm², whereas in auxin-treated DDM6 worms, the area ranged from 0.64 µm² to 2.68 µm², with an average size of 1.18 µm² (Figure 3d). Dietary OA supplementation led to a partial recovery of total fat storages and restored lipid droplet size (Figure 3b, d).

We have recently demonstrated that multidimensional NMR spectroscopy, combined with uniform ^13^C-isotopic enrichment of *C. elegans*, enabled the dissection of lipid composition in LDs of live worms. (Hernández-Cravero et al., 2024). Lipid analysis of live, ^13^C-isotopically enriched, auxin treated animals confirmed a reduction in UFA, PUFA, and TAG (Figure 3e and Figure S4a), consistent with the decreased fat deposits and reduced LD size observed via fluorescence microscopy. NMR analysis of the supernatants confirmed that lipid signals originated from molecules within the intact worms (Figure S4b). Exogenous supplementation with non-isotopically labelled OA restored TAG levels but not UFA or PUFA levels as described in more detail below. These observations were confirmed by complementary NMR experiments on total lipid extracts from the ^13^C-isotopically enriched worms (Figure 3f). As mentioned above, upon OA supplementation of auxin treated animals, TAG levels were restored, but PUFA levels were approximately 90% lower than those in ethanol-treated control animals and 40% lower than in auxin-treated animals without OA supplementation. This seemingly opposing behavior is explained by the fact that only endogenous TAG and fatty acyl chains that were generated during the growth on isotopically enriched bacteria are uniformly ^13^C-enriched, and thus visible by NMR (Hernández-Cravero et al., 2024). The reduction in UFA and TAG levels observed in the presence of auxin thus directly reflects the decrease in these lipid species within the animals. In contrast, exogenously added OA is not isotopically enriched and thus remains NMR-invisible. As a result, newly synthesized lipid molecules that incorporate exogenous OA, such as UFA and PUFA fatty acyl chains, will not be detected in ^13^C correlation spectra (Figure S5). Therefore, the reduction in the PUFA signal following auxin/OA treatment, compared to auxin alone, suggests that the turnover of endogenous ^13^C-enriched PUFAs increases in the presence of OA, with new PUFAs being synthesized using exogenous OA as a substrate. This observation aligns with the lack of FAT-7 activity in the presence of auxin.

## Discussion

In the present work, we have introduced a novel *C. elegans* strain that enables conditional depletion of UFA. Such strain, allows for quick decrease of all sources of UFA, with a simple intervention: placing the worms on auxin-supplemented media at the desired larval stage. We successfully replicated the lethal phenotype of the *fat-6; fat-7 fat-5* triple mutant (Brock et al. 2006) in DDM6 through auxin treatment, demonstrating the efficiency of depleting AID-tagged FAT-7. Moreover, we showed that the observed phenotypes upon auxin treatment were specific to the loss of Δ9 desaturase activity as they were rescued by supplementing the media with OA, the product of this enzyme. Moreover, GC-MS and NMR analysis on lipid extracts (Table 1 and Figure 2e) revealed that the reduction in UFA content when worms reached the one-day adult stage is due to a decrease in PUFA levels rather than MUFA content. We have also shown that longer exposure to auxin does not cause significant changes in the SFA/UFA ratio (Table S1, Figure 2d).

High-resolution *in vivo* NMR analysis of DDM6 worms treated with auxin confirmed the absence of FAT-7 activity and a decrease in UFA and PUFA levels. Additionally, TAG content in LDs was lower in auxin-treated worms, further supporting the role of FAT-7 in regulating TAG biosynthesis and its incorporation into LDs (Shi et al. 2013; Wang et al. 2022; Han et al. 2024). OA supplementation restored TAG and UFA levels, as demonstrated by GC-MS and ^1^H NMR spectroscopy, and also replenished adiposity and LD content. Heteronuclear ^1^H-^13^C correlation NMR experiments revealed that the replenishment of fatty acyl chains in TAG molecules originated from exogenously added OA. Furthermore, OA addition resulted in a marked decrease of ^13^C-isotopically enriched PUFAs, which had been synthesized *de novo* by the worms, indicating an increase in lipid turnover stimulated by OA. These findings align with previous research showing that RNAi depletion of FAT-6 and FAT-7, which reduces OA and UFA production, significantly diminishes the incorporation rate of most fatty acids into membrane lipids (Dancy et al. 2015). Intriguingly, OA has been shown to accelerate β-oxidation of lipids through a Sirt- 1/PGC1α-dependent mechanism in skeletal muscle cells (Lim et al. 2013), suggesting that the effects of OA on lipid turnover may be widespread among higher organisms. Our results supports the use of NMR for monitoring lipid turnover (Lin et al. 2021; Lin et al. 2024) and highlights our system as an experimental model to further explore this mechanism, which remains poorly defined (Perez and Van Gilst 2008; Watts and Ristow 2017).

The depletion of total UFA synthesis at the L3 stage showed noticeable phenotypes in DDM6 strain. These animals had dramatically reduced survival at low temperature (Figure 2a) and reduced brood size (Figure 2b). During oogenesis, the developing germ cells need a constant supply of resources to keep growing. To do so, somatic resources are transferred to the germline through the action of vitellogenins, which assemble and transport lipids as yolk from the intestine to the developing oocytes (Schneider 1996; Hall et al. 1999; Lynn et al. 2015). The transport of yolk to the oocyte relies on proper fatty acid composition (Chen et al. 2016), and conditional depletion of total PUFA synthesis by auxin exposure could accumulate excess yolk material in the pseudocoelom leading to reduced brood sizes (Figure 2b).

Previous work showed that a *fat-6; fat-7 fat-5* triple mutant can grow when supplemented with OA, LA, and EPA (Brock et al. 2006). However, in that context, when L3/L4 larvae were transferred from supplemented to unsupplemented plates they became thin, sterile adults that died early (Brock et al. 2006). In contrast, DDM6 L3 larvae previously grown on control plates and then exposed to auxin remain viable and fertile, albeit with a smaller brood size, and their eggs develop into fertile adults on auxin-free plates (data not shown). Thus, UFAs synthesized before FAT-7 degradation are more effective than exogenous fatty acid supplementation in supporting development into fertile adults. This unique property of the AID system in DDM6 can be leveraged for future studies of germline defects induced by complete depletion of de novo UFA biosynthesis.

Finally, we identified a critical role for endogenous OA (OA) production in facilitating the transition from the L2 larval stage to adulthood (Figure 1b, c). Interestingly, *fasn-1* RNA knockdown also leads to developmental arrest of *C. elegans* at the L2 stage (Li and Paik 2011). FASN is well known for its role in the biosynthesis of palmitate (16:0), a key building block of UFAs (Figure 1a). However, a recent report demonstrated that beyond its essential role in fatty acid synthesis, the C-terminal fragment of FASN-1 (FAS-CTF) also has a regulatory role in mitigating stress, which cannot be rescued by palmitate (Wei et al. 2024). Therefore, it is possible that, in addition to its structural role, oleate or PUFAs—rather than stearoyl-CoA desaturase (SCD) alone—may have a regulatory function in the L2-to-L3 transition. However, the underlying mechanism remains unknown. One possible explanation is that OA or certain OA-derived molecules might act as intermediates capable of activating signaling pathways related to growth. In this context, previous studies have demonstrated the role of UFAs in modulating the activity of TRPV channels through sensory transduction (Kahn-Kirby et al. 2004) and the action of endocannabinoids (UFA-derived molecules) in regulating energy metabolism in *C. elegans* (Galles et al. 2018; Hernandez-Cravero et al. 2022).

In conclusion, we used the auxin-inducible degradation system targeting the *fat-7* gene in a double mutant (*fat-6, fat-5*) to develop a method that externally induces OA auxotrophy in *C. elegans*. We show that auxin exposure during larval development conditionally halts *de novo* total UFA synthesis, which can be reversed upon cessation of auxin exposure or OA supplementation. This system is valuable for studying processes involving membrane remodeling (e.g., fluidity, curvature, vesicle and protein trafficking or signaling) and germline maintenance. Unlike RNAi or double mutants, it avoids pleiotropy and is less labor-intensive, providing significant advantages for future studies.

## Acknowledgments

We acknowledge Cecilia Vranych for assistance with *C. elegans* growth and maintenance

